# Protease-independent production of poliovirus virus-like particles in *Pichia pastoris*: Implications for efficient vaccine development and insights into capsid assembly

**DOI:** 10.1101/2022.09.16.508263

**Authors:** Lee Sherry, Jessica J. Swanson, Keith Grehan, Huijun Xu, Mai Uchida, Ian M. Jones, Nicola J. Stonehouse, David J. Rowlands

## Abstract

The production of enterovirus virus-like particles (VLPs) which lack the viral genome have great potential as vaccines for a number of diseases, such as poliomyelitis and hand, foot-and-mouth disease. These VLPs can mimic empty capsids, which are antigenically indistinguishable from mature virions, produced naturally during viral infection. Both in infection and in vitro, capsids and VLPs are generated by the cleavage of the P1 precursor protein by a viral protease. Here, using a stabilised poliovirus 1 (PV-1) P1 sequence as an exemplar, we show the production of PV-1 VLPs in *Pichia pastoris* in the absence of the potentially cytotoxic protease, 3CD, instead using the porcine teschovirus 2A (P2A) peptide sequence to terminate translation between individual capsid proteins. We compare this to protease-dependent production of PV-1 VLPs. Analysis of all permutations of the order of the capsid protein sequences revealed that only VP3 could be tagged with P2A and maintain native antigenicity. Transmission electron microscopy of these VLPs reveals the classic picornaviral icosahedral structure. Furthermore, these particles were thermostable above 37°C, demonstrating their potential as next generation vaccine candidates for PV. Finally, we believe the demonstration that native antigenic VLPs can be produced using protease-independent methods opens the possibility for future enteroviral vaccines to take advantage of recent vaccine technological advances, such as adenovirus-vectored vaccines and mRNA vaccines, circumventing the potential problems of cytotoxicity associated with 3CD, allowing for the production of immunogenic enterovirus VLPs *in vivo*.

## Introduction

Poliomyelitis, a devastating paralytic and potentially fatal disease caused by poliovirus (PV) has been responsible for many global epidemics over the past century. The introduction of the Global Polio Eradication Initiative (GPEI) in 1988, has resulted in >99% reduction in paralytic poliomyelitis cases globally and the goal of the initiative is to eliminate the virus (1). This reduction in paralytic PV cases has resulted from the widespread application of two vaccines; the live-attenuated oral PV vaccine (OPV) and the inactivated PV vaccine (IPV) (2). Vaccine deployment has also led to wild-type (wt) PV-2 and wt PV-3 being declared eradicated in 2015 and 2019, respectively, with only wt PV-1 still circulating in Afghanistan and Pakistan (3). As we near eradication, biosafety concerns surrounding both OPV and IPV as each requires large scale virus growth and thus has the potential to re-introduce the virus into the environment.

Whilst IPV provides excellent humoral immunity which protects against paralytic disease, it induces little, if any, mucosal immunity and does not prevent viral replication in the gut, thus allowing virus transmission within a population (4). The contribution of OPV to the near-eradication of PV is immense, although control of polio by IPV alone has been achieved in some regions. However, the attenuated virus can quickly revert to virulence, leading to vaccine-associated paralytic poliomyelitis (VAPP) and, in areas with low vaccine coverage, the spread of vaccine-derived PV (cVDPV) (5). Unfortunately, cVDPV cases now outnumber wt PV cases globally (1). Additionally, OPV can recombine with other PVs or PV-like enteroviruses to gain virulence. This, alongside chronic shedding of VDPV into the environment from immunocompromised individuals, highlights the risks associated with the continued use of OPV for eradication of polio (6, 7). However, the recent introduction of nOPV2 vaccine, which significantly decreases the likelihood of reversion and recombination, may help to reduce VDPV circulation and therefore assist in PV eradication (8).

PV belongs to species *Enterovirus C* within the picornavirus family of positive-sense RNA viruses and has a 7.5 kb genome. The majority of the genome encodes a large continuous open-reading frame (ORF), which is proteolytically-processed after translation into mature protein products. In addition, a short upstream ORF (uORF) has recently been identified and shown to be important in *ex vivo* organoid infection by Echovirus 7 (9). The major ORF comprises 3 distinct regions; P1, which encodes the viral capsid proteins, and P2 and P3, which encode the non-structural proteins. Proteolytic cleavage of the polypeptide by the viral proteases, 2A^pro^, 3C^pro^ and 3CD, occurs co- and post-translationally yielding the mature viral proteins required for viral replication (10). The viral protease precursor, 3CD, specifically cleaves P1 into the individual capsids proteins, VP0, VP3 and VP1 (11, 12). A further maturation cleavage of VP0 into VP4 and VP2 is associated with encapsidation of viral RNA and results in enhanced particle stability (13, 14).

The mature PV virion is a ~30 nm icosahedral capsid, comprised of 60 copies of VP1-VP4, containing the viral genome (15). During infection, particles lacking viral genome are also produced. These particles, termed empty capsids (ECs), are antigenically indistinguishable from mature virions, although VP0 remains uncleaved (16). Recombinant ECs have potential as virus-like particle (VLP) vaccines, however wt ECs are inherently unstable and their antigenic conformation changes at lower temperatures than is the case for mature viral particles (16, 17). This conformational change results in a minor expansion of the particles, but has significant consequences for their antigenicity. ECs readily convert from the native antigenic form (termed D-Ag) to the non-native form (termed C-Ag). Although the D-antigenic form induces protective immune responses to PV, the C-antigenic form does not (17–19). Therefore, recombinant VLP vaccines against PV must retain the D-antigenic conformation.

VLPs are a safe and attractive option as recombinant vaccines as they mimic the repetitive structures of virions but lack the genome and are non-infectious. Both the hepatitis B virus (HBV) and human papillomavirus (HPV) vaccines are licensed VLP-based vaccines produced in yeast and insect cells, respectively (20–22). Following the initial demonstration of the expression of PV VLPs in yeast in 1997 (23), these have been produced in several systems in recent years, including mammalian, plant and insect cells (24–30). However, most of these systems rely on the co-expression of the viral protease, 3CD, to cleave the viral structural protein to produce VLPs. There are some concerns around the potential cytotoxicity of 3CD and its potential impact on VLP yield in recombinant systems. Additionally, 3CD has been shown to induce apoptosis in mammalian cells (31), which may limit the application of newly licensed vaccine technologies such as viral vectors and mRNA vaccine technologies to produce next-generation poliovirus vaccines.

We have previously demonstrated the production of PV VLPs using *Pichia pastoris*, and modulated the expression of 3CD through a number of molecular approaches (30). Interestingly, Xu et al. reported an insect cell expression system which yielded PV VLPs in the absence of the viral protease by splitting the P1 precursor protein across 2 ORFs, with VP3 and VP0 under the control of one promoter and separated by the porcine teschovirus 2A peptide (P2A), and VP1 under the control of a second promoter (28), which also changes the natural order of proteins on the P1 precursor, i.e. VP0-VP3-VP1.

Here, we investigated the potential for protease-independent VLP production in *Pichia pastoris*. We show that stabilised PV-1 VLPs (32) can be produced in *Pichia* without the viral protease, 3CD, using instead the P2A peptide sequence to terminate translation between individual capsid proteins, and compare this to protease-dependent production of PV-1 VLPs. Analysis of all permutations revealed that only VP3 could be tagged with P2A and maintain native antigenicity. Transmission electron microscopy of these VLPs reveals their classic picornaviral icosahedral structure. Finally, we show that these particles are thermostable above 37°C, demonstrating their potential as next-generation polio vaccine candidates.

## Results

### Protease-independent production of poliovirus structural proteins

Previously, we reported the production of PV VLPs in *Pichia Pastoris* using dual promoter constructs to separately express P1 and the viral protease, 3CD. However, the over-expression of 3CD can be cytotoxic and may reduce the yield of PV VLPs in heterologous systems (30, 31). Xu et al. demonstrated production of protease-independent PV VLPs in native conformation in insect cells using recombinant baculoviruses to express the viral structural proteins under the control of two separate promoters, with VP3 and VP0 separated by P2A transcribed from the first promoter and VP1 under the control of the second promoter (Xu et al. 2019). 2A is a short peptide sequence present at the C terminus of the P1 region of aphthoviruses which interrupts translation to separate the P1 and P2 regions without proteolytic cleavage. Here, we undertook a detailed study of the protease-independent expression of PV-1 VLPs in *Pichia* using a matrix of constructs based on a PV-1 sequence which includes several stabilising mutations (32) (Fig. 1A). We compared protein expression and VLP assembly from these constructs with the dual alcohol oxidase (AOX) promotor P1/3CD system we reported previously, together with a modified construct in which the VP1 sequence was extended at the C-terminus with a 6xHIS tag.

**Figure 1:**
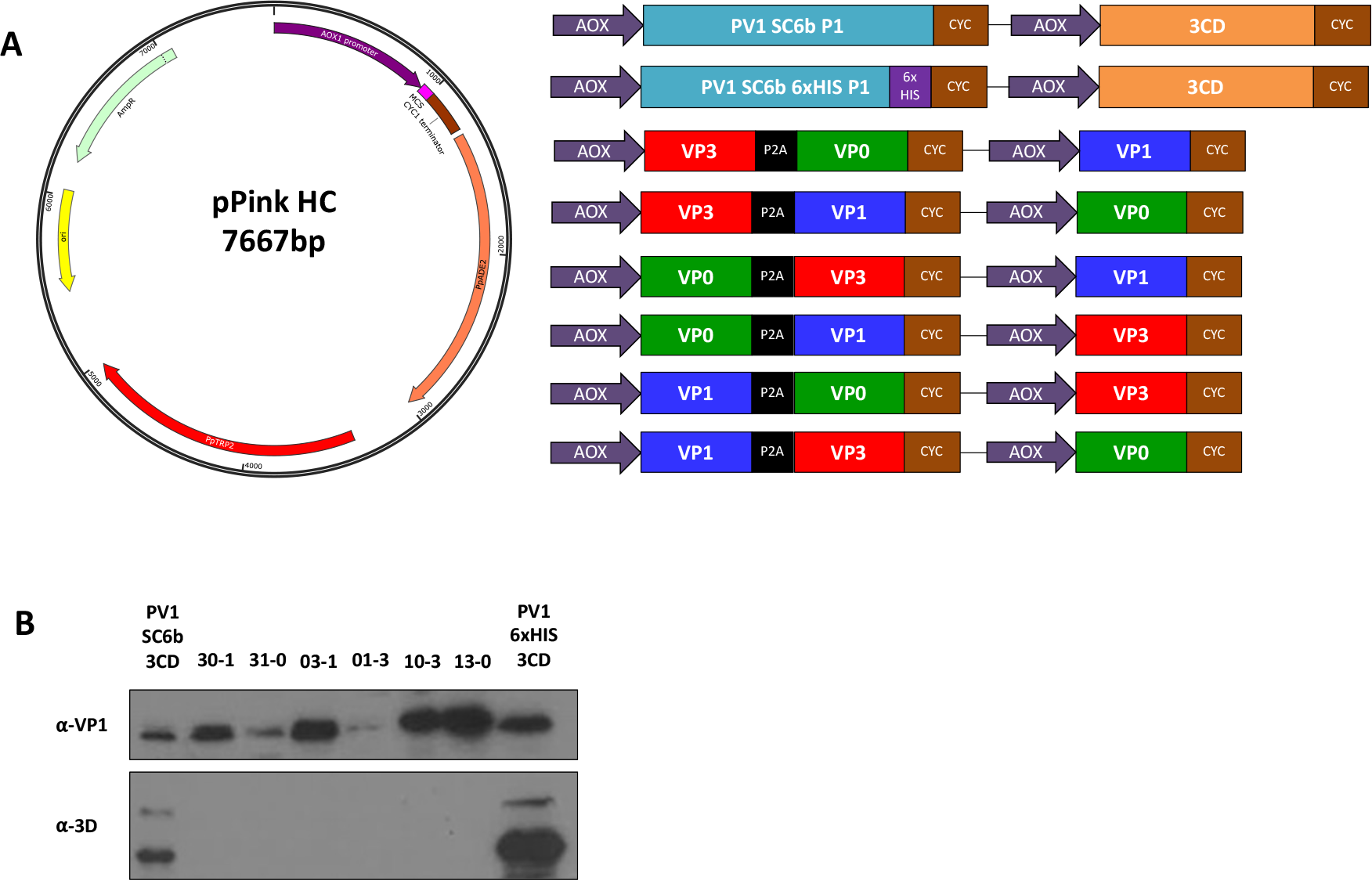
Schematic of expression cassettes and immunblots. **A**: The pPink HC expression vector with the dual alcohol oxidase (AOX) promoter expression constructs which drive the expression of the P1 capsid protein and the viral protease, 3CD or the expression of protease-free constructs with VP0 (green), VP3 (red), and VP1 (blue). **B**: Immunoblot for PV VP1 and 3D. Samples were collected 48 hours post-induction and lysed using 0.1 M NaOH. All samples were boiled in 2x Laemlli buffer and separated by SDS PAGE prior to analysis by immunoblot using mouse monoclonal α-VP1 and rabbit monoclonal α-3D antibody. The figure is a representative example of three separate experiments for each construct.

*Pichia* colonies, transformed with each construct, were tested for the level of expression. Samples from small-scale expression were collected 48 hours post-induction and analysed for correct processing, either by 3CD or P2A, using an anti-VP1 antibody (Fig. 1B). All constructs produced VP1, albeit at different levels. Both protease-containing constructs, PV-1 SC6b P1 and PV-1 SC6b 6xHIS, and the protease-independent constructs in which VP1 was upstream of 2A or was produced from a separate promotor, produced readily detectable levels of VP1. However, in constructs where VP1 followed the P2A peptide, VP3-2A-VP1 VP0 and VP0-2A-VP1 VP3 respectively, the level of VP1 was markedly lower. This suggests that, although in an optimum context P2A can facilitate a 1:1 ratio of up- and down-stream protein expression in mammalian cells, the surrounding sequence or the re-initiation event following 2A-mediated interruption of translation may influence the translation rates in *Pichia*. As expected, anti-3D reactivity was only seen with lysates of the protease-containing constructs, where two major bands were detected at approximately 72 and 55 kDa respectively (Fig. 1B).

### VLPs produced from protease-dependent and -independent constructs sediment similarly

To determine VLP assembly competence of the protease-independent constructs, high-expressing *Pichia* clones of each were cultured to high-density and expression induced with 0.5% methanol (v/v) and cell pellets collected 48 hours post-induction. The resuspended pellets were homogenised at ~275 MPa and the resultant lysates purified through chemical precipitation and differential centrifugation steps culminating in 15-45% sucrose gradients. With the exception of the VP0-2A-VP3 VP1 construct, immunoblot analysis of gradient fractions detected VP1 in fractions consistent with the presence of VLPs (Fig. 2) with intensities that were broadly consistent with the total protein analysis (Fig. 1). Constructs in which 2A followed VP0 produced lower yields of VLPs suggesting a conformation incompatible with efficient assembly. Minor differences in peak fractions was also noted with those constructs containing VP1-2A or VP1-His trending to peak higher in the gradient.

**Figure 2:**
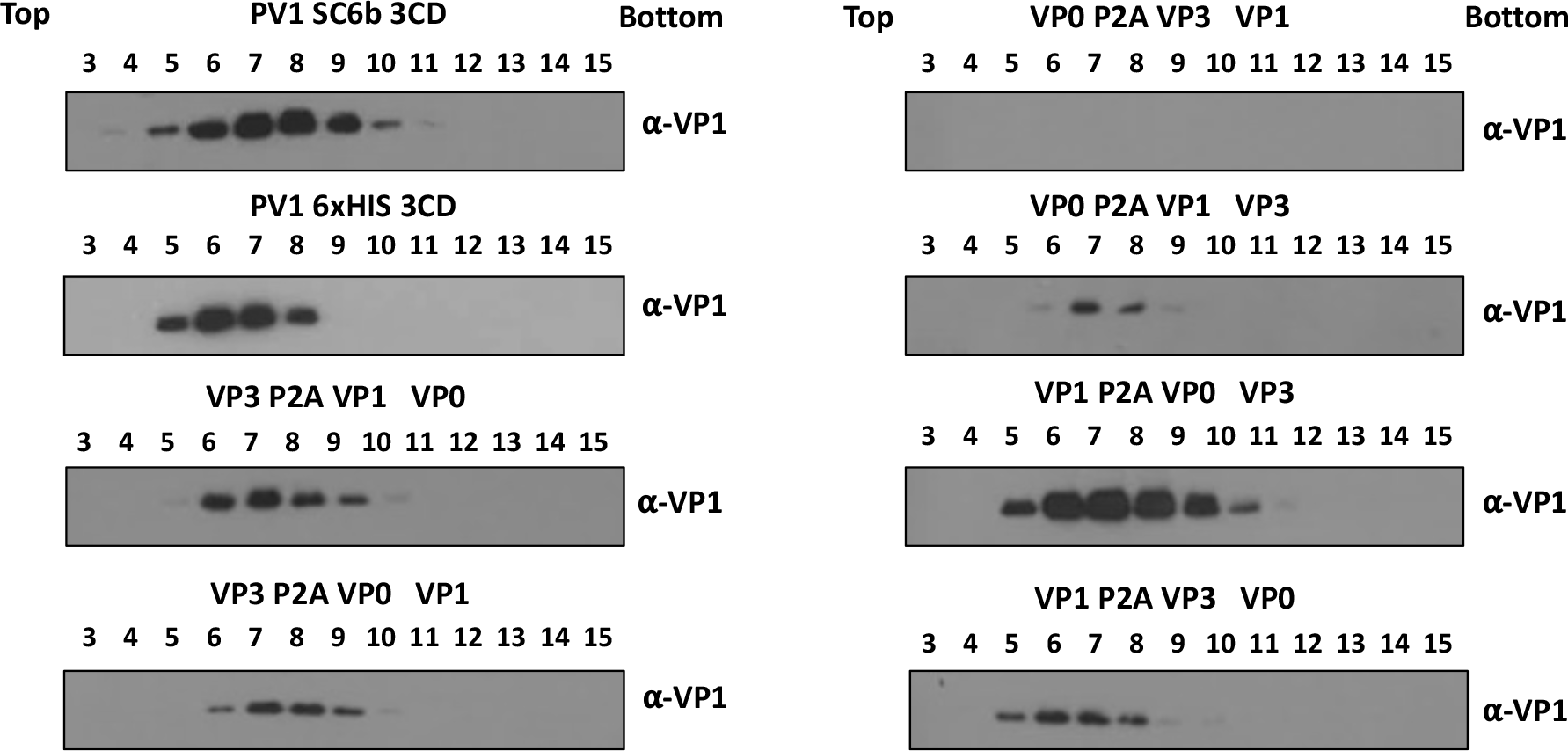
Gradient purification of protease-dependent and protease-free PV-1 SC6b VLPs. All samples were purified by ultracentrifugation and fractioned from top to bottom in 1 mL aliquots. A 12 μL sample from each fraction was then taken and boiled in 5x Laemmli buffer and separated by SDS PAGE prior to analysis by immunoblot using mouse monoclonal α-VP1. The figure is a representative example of three separate experiments for each construct.

### VP3-P2A VLPs are D-antigenic

To confirm the antigenicity of the VLPs, the peak fractions were analysed by enzyme-linked immunosorbent assay (ELISA) using a standard protocol established by the National Institute for Biological Standards and Control (NIBSC) with the current inactivated vaccine (BRP) as a positive control (32). As expected, the PV-1 SC6 3CD produced both D and C antigenic VLPs, with the ratio of D:C largely in favour of D-antigenic particles (Fig 3). Intriguingly, the addition of a VP1 C-terminal 6xHIS tag on this construct reversed the ratio, with the majority of particles found in the C-antigenic conformation. This was also true for both protease-independent VP1-P2A constructs, only C-antigenic particles were detected, suggesting that the addition of a C-terminal tag to VP1 leads to assembly of particles, which readily convert to C-antigenicity, or never achieve the D-antigenic conformation despite the presence of stabilising mutations (Fig 3).

**Figure 3:**
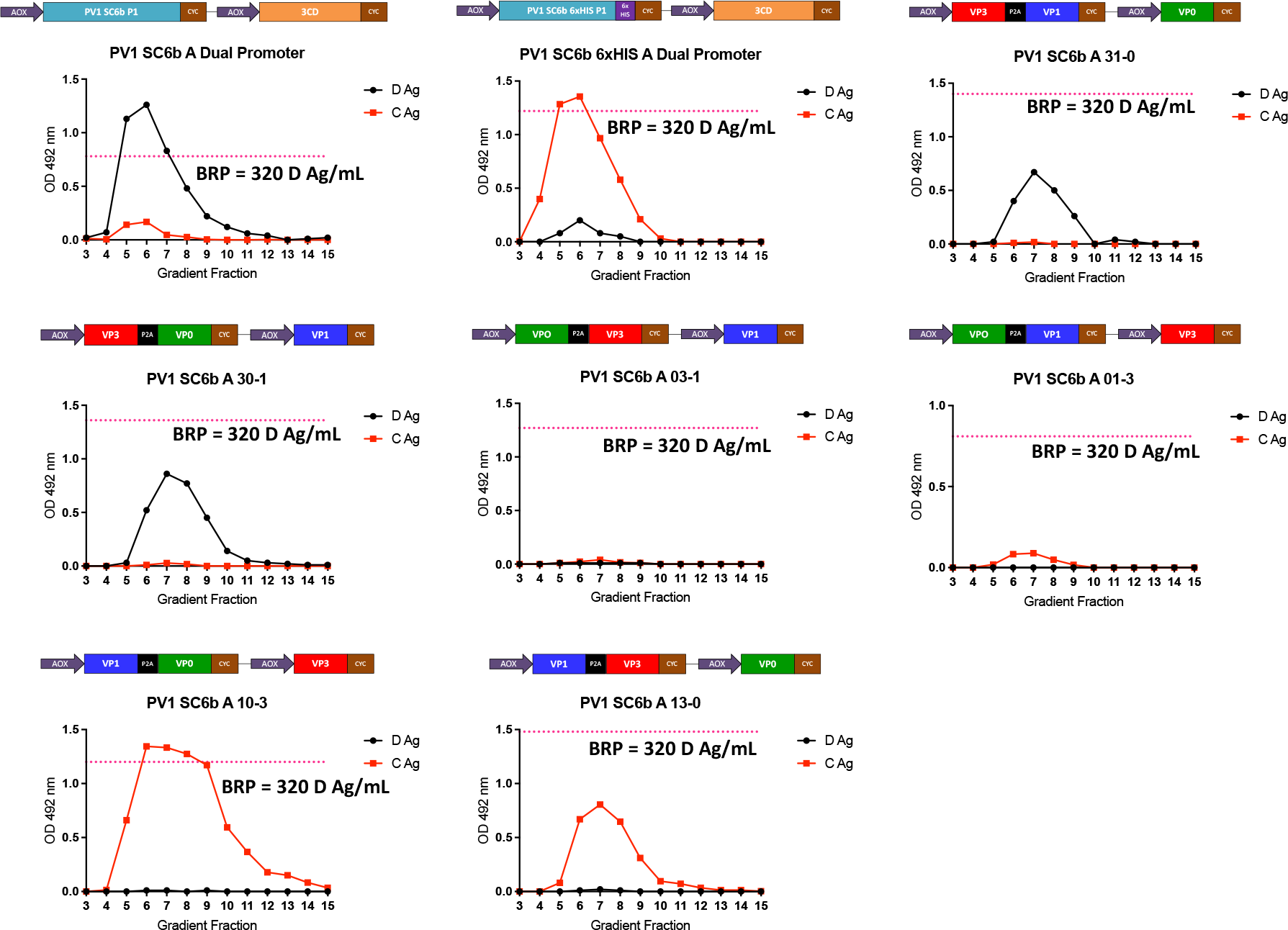
Antigenicity of PV-1 VLPs. Reactivity of gradient fractions with Mab 234 (D antigen) and Mab 1588 (C antigen) in ELISA. The pink dashed line represents the positive control, BRP, for the D antigen ELISA. OD at λ=492 nm is represented in arbitrary units. The figure is a representative example of three separate experiments for each construct.

Despite low level VP1 expression in the lysates of the VP0-2A-VP1 VP3 construct (Fig 2), no VLPs were detected in the gradient fractions (Fig 3) and although strong VP1 expression was apparent for the VP0-2A-VP3 VP1 construct (Fig 2) only low levels of assembled VLPs, which were entirely in the C-antigenic conformation, were observed (Fig 3).

Interestingly, both VP3-P2A constructs produced D-antigen reactive particles with little to no detectable C-antigen, suggesting that addition of a C-terminal tag to VP3 is not detrimental to the antigenicity of assembled particles, consistent with the data previously reported following expression in insect cells (Xu et al. 2019). These data demonstrate that D-antigenic VLPs can be produced in *Pichia* without the requirement for co-expression of the viral protease, 3CD.

To further characterise the assembled VLPs, peak fractions of the productive constructs were pooled and concentrated using 100 kDa centrifugal concentrators and assessed by immunoblot and ELISA (Fig. 4A and 4B). The concentrated VLPs were interrogated by immunoblot using a number of different antibodies (Fig. 4A). As expected, the relative levels of VP1 and VP0 were similar for each VLP and only the PV1 SC6b 6xHIS showed reactivity to the anti-HIS antibody. The anti-2A antibody detected the tagged proteins VP3-P2A and VP1-P2A (~27 kDa and ~33 kDa, respectively) as expected. However, for the VP3-P2A-VP0 VP1 VLPs, the VP3-P2A band was weaker than that seen with VLPs derived from the VP3-2A-VP1 VP0 construct. This may be a consequence of enhanced proteolytic degradation of the 2A tag in the context of the VP3-P2A-VP0 VP1 VLPs.

**Figure 4:**
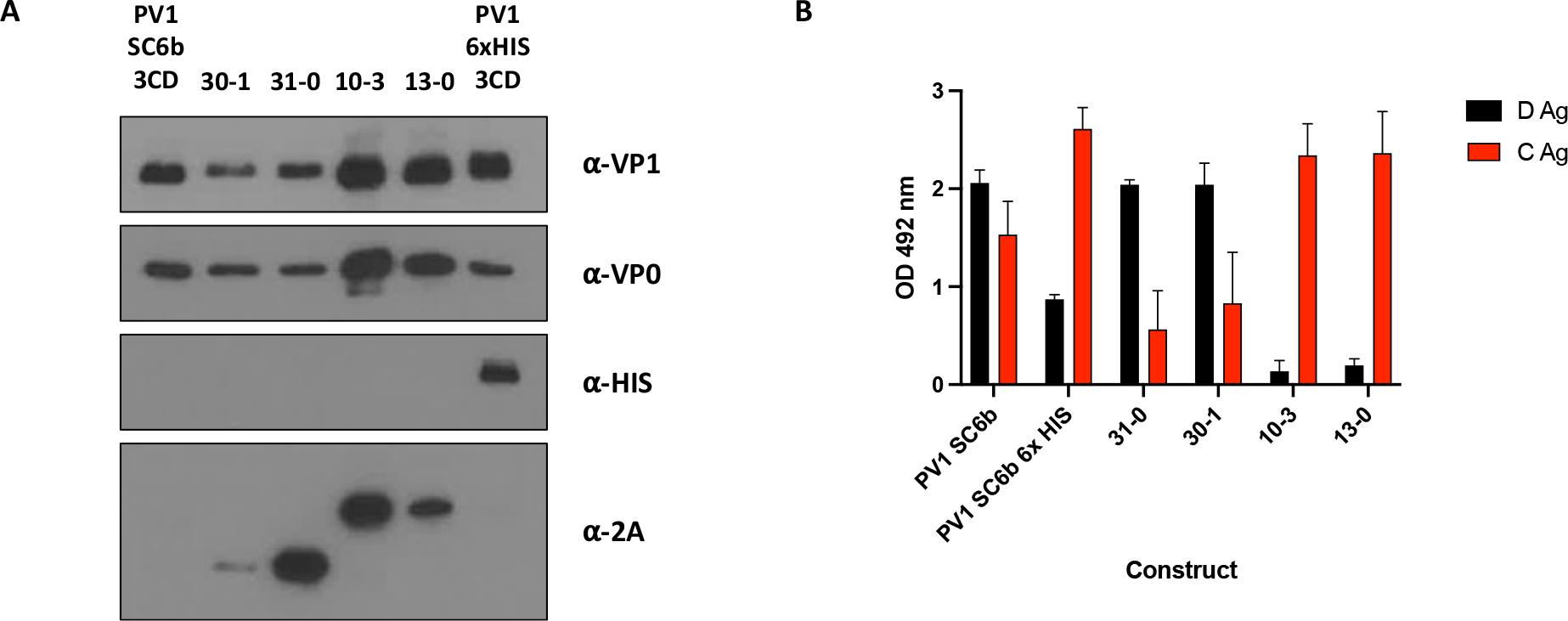
Characterisation of protease-dependent and protease-free PV-1 SC6b VLPs. **A:** Immunoblot for PV VP1, VP0, 6xHIS and 2A peptide. Peak gradient fractions for each VLP preparation were concentrated using 100 kDa microcentrifuge concentrators (Amicon) to ~100 uL. All samples were mixed 1:1 with 2x Laemlli buffer, boiled and separated by SDS PAGE prior to analysis by immunoblot using mouse monoclonal α-VP1, a rabbit polyclonal α-VP0, mouse monoclonal α-HIS and mouse monoclonal α-2A. **B:** Antigenicity of concentrated PV-1 VLPs. Reactivity of concentrated fractions with Mab 234 (D antigen) and Mab 1588 (C antigen) in ELISA. OD at λ=492 nm is represented in arbitrary units. The figure is a representative example of three separate experiments for each construct.

We examined the C- and D-antigenic composition of the VLPs in more detail by ELISA following their concentration from the sucrose gradient fractions (Fig. 4B). Both D- and C-antigenic forms were present in VLPs derived from protease-containing constructs, although the D:C ratio was greatly reduced in the P1-6xHIS tagged construct. Little or no D-antigen was detected in the concentrated VP1-P2A VLPs from the VP1-P2A constructs, further indicating that tagged VP1 results in the assembly of C antigenic VLPs. In contrast, the VLPs including VP3-2A produced high levels of D-antigenic material and low levels of C-reactivity.

### Protease-independent VLPs maintain classic picornavirus morphology and are thermally stable at physiologically relevant temperatures

Figure 5A shows representative negative-stain EM images of concentrated protease-independent VLPs and the protease-dependent VLPs. Particles of ~30 nm diameter with typical VLP morphology were seen in all samples, consistent with previous EM images of poliovirus virions and empty capsids and *Pichia*-derived PV-1 VLPs (30). Smaller particles were also seen, especially in the protease-independent samples, which are likely to be *Pichia* alcohol oxidase or fatty acid synthetase which have been shown previously to co-purify with VLPs (33).

**Figure 5:**
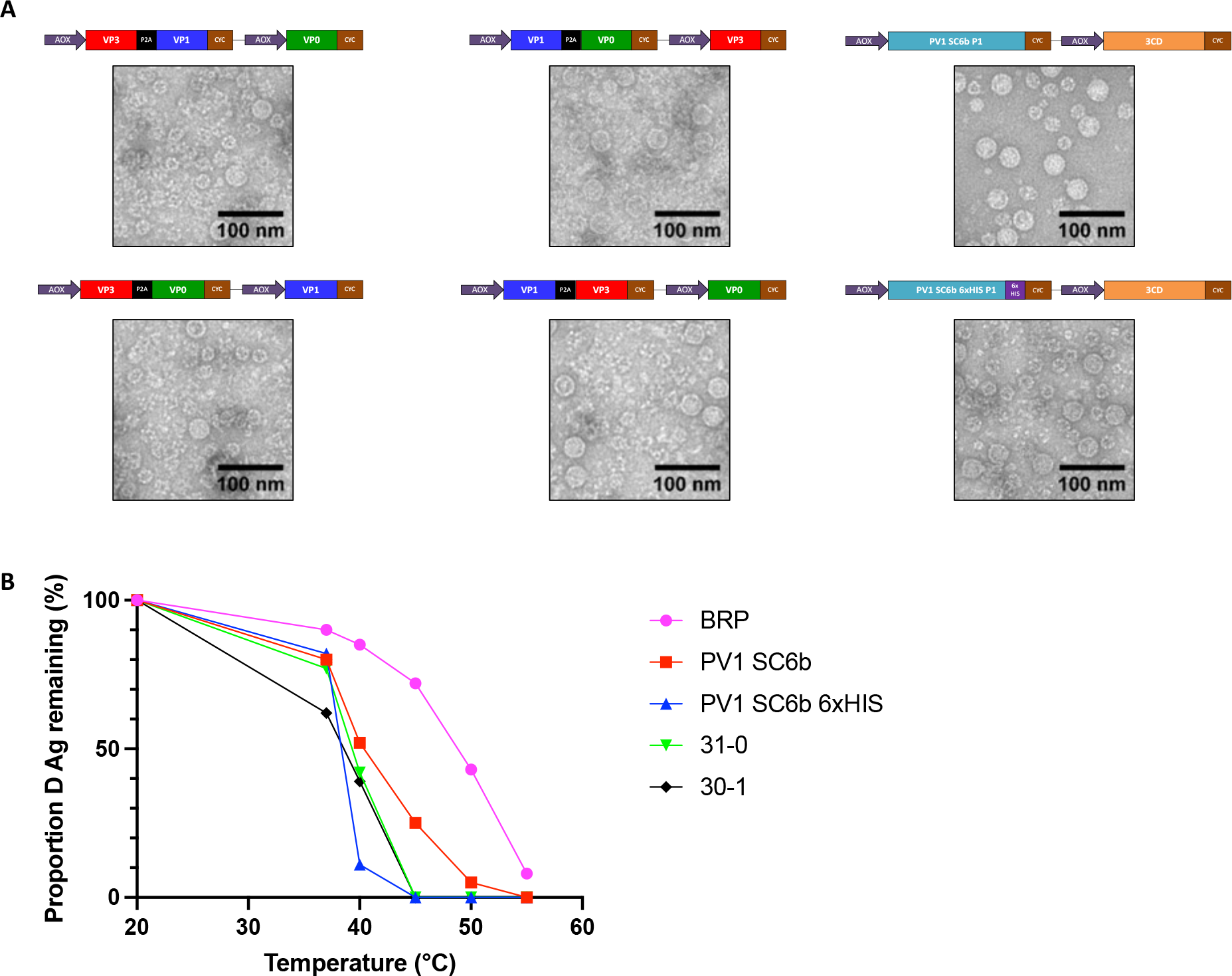
Transmission Electron Microscopy of VLPs and Thermostability. **A:** Representative micrographs of protease-dependent and protease-free PV-1 SC6b VLPs. (Scale bar shows 100 nm). **B:** Reactivity of purified PV-1 SC6b VLPs and BRP aliquots to D-antigen specific MAb 234 in ELISA after incubation at different temperatures for 10 minutes, normalised to corresponding aliquot incubated at 4 °C. The figure is a representative example of two separate experiments for each construct

Potential vaccines will need to withstand physiologically relevant temperatures and so the D-antigenicity of VP3-P2A containing VLPs, protease-derived VLPs, and BRP, was assessed following exposure to increasing temperatures (Fig. 5B). The positive control, BRP, was the most thermally stable only showing a 50% decrease in D antigenicity at ~50°C whereas PV-1 SC6b 3CD VLPs lost 50% D Ag ~41°C, although VLPs with VP1 6xHIS were least stable with D Ag dropping sharply above 37°C and retaining only 11% D-antigenicity at 40°C. Importantly, both VP3-P2A containing VLPs maintained D-antigenicity above 37°C (62% and 77% for VP3-P2A-VP0 VP1 VLPs and VP3-P2A-VP1 VP0 VLPs, respectively). These results suggest that protease-free production of PV VLPs can yield particles which are antigenically stable at physiologically relevant temperatures, suggesting the potential to produce a long-lasting immune protection against PV.

## Discussion

The current PV vaccines require the production of large amounts of infectious virus, with potential to reintroduce the virus to the environment (2, 4). Moreover, continued use of OPV has contributed to the global increase in cVDPV cases, which now exceed those caused by wt PV (7). However, these vaccines are hugely important for eradication of PV, especially in light of the recently licensed nOPV2, which has been intelligently designed to reduce the likelihood of the two major concerns surrounding OPV, reversion and recombination. (8). With this newly available vaccine, and advances with improved PV-1 and PV-3 OPV vaccines, there is increased optimism that a polio-free world is achievable. However, for the complete elimination of polio alternative PV vaccines which do not require the growth of large quantities of infectious virus for their production still require investigation. VLP vaccines produced using a heterologous expression system may address this requirement. This approach has been successful in a number of expression systems, including plant, insect cell, MVA-based mammalian cell expression systems and, as we show here, yeast systems such as *Pichia Pastoris* (24–30, 34). However, the 3C protease, the active component of the P1 cleavage specific precursor 3CD, has been shown to have a toxic effect on cell viability. Whilst 3CD should be less toxic than 3C, it has the potential for cellular toxicity (12, 31), which may reduce the yield of VLPs produced in heterologous expression systems, as highlighted by our failure to select viable colonies in *Pichia pastoris* when expressing 3CD under a constitutive promoter. However, a recent publication indicated the potential of protease-independent production of PV VLPs using an insect cell expression system in which the two structural proteins, VP0 and VP3, were separated by inserting a P2A peptide, and with VP1 under the control of a second promoter (28). Here, we explored all permutations of P2A containing protease-independent VLP constructs using *Pichia pastoris* as a heterologous expression system.

We investigated the efficiency of viral capsid protein expression by immunoblot analysis (Fig 1B.) Good levels of VP1 were produced from both protease-dependent constructs and from each protease-independent construct where the VP1 ORF was immediately after the promoter, with or without the P2A peptide. However, lower levels of VP1 were seen when the VP1 ORF was placed after the P2A peptide. This suggests that although in the appropriate context the P2A peptide is highly efficient at re-initiating translation following pausing in mammalian cells, the efficiency of this re-initiation event in *Pichia* is lower (35). This also suggests that the individual structural proteins are unlikely to be at equimolar amounts within the cell, which would inevitably reduce capsid assembly efficiency. Therefore, there is potential to improve this process in future either by selection of an alternative 2A peptide or by the expressing all three structural proteins individually from different promoters (36).

Following conformation that each of these constructs produced the viral structural proteins, we compared the production of VLPs from the protease-independent constructs with those from the protease-dependent constructs, which were previously shown to produce VLPs in *Pichia* (30). Interestingly, almost all of the protease-independent constructs produced material, which sedimented in gradients similarly to VLPs produced from the protease-dependent constructs. The exceptions were VP0-P2A-VP3 VP1 and the VP0-P2A-VP1 VP3 constructs which produced less material than other protease-dependent and protease-independent constructs (Fig 2.) Using conformation-specific antibodies in ELISAs of sucrose gradient fractions we saw no signal from the VP0-P2A-VP3 VP1 construct as expected, and only low levels of C-antigenic and no D-antigenic reactivity from the VP0-P2A-VP1 VP3 construct. These results suggest that addition of the P2A peptide to the C-terminal end of VP0 has detrimental effects on particle assembly.

Whilst VP1-P2A tagged constructs produced assembled VLPs, as demonstrated by sucrose gradient centrifugation analyses (Fig 2.) and morphologically by TEM (Fig 5A.), these were entirely in the C-antigenic conformation (Fig 3 & 4A). This dominance of the C-antigenic conformation was also seen when a C-terminal 6xHIS tag added to VP1 in the protease-dependent construct, although some D-antigenicity was detected. Virus particles with 2A peptide still attached have been previously described for foot-and-mouth disease virus, another member of the picornavirus family, although the particle expansion and antigenic conformational changes seen with PV are not observed here (37). However, in closer relatives to PV, VP1 C-terminal extensions which include a motif integral to viral entry process have been found in Coxsackieviruses, echoviruses and human parechoviruses (38–40).

Interestingly, both VP3-P2A constructs produced D-antigenic VLPs, in agreement with previous work in insect cells which showed that a dual promoter construct containing VP3-P2A-VP0 under the control of one promoter and VP1 under the control of a second promoter produced D-antigenic VLPs (28). Our initial sucrose gradient analyses (Fig 2.) suggested that these constructs produced little to no detectable C-antigenicity, whilst maintaining good levels of D-antigenicity. C-antigenicity was detectable after concentration of the VLPs, although at a much lower level than D Ag and both constructs produced similar D:C ratios (Fig 4B.)

Intriguingly, when we assessed the levels of P2A peptide in the concentrated VLP samples, the VP3-P2A-VP0 VP1 VLPs showed less reactivity to the 2A antibody than the VP3-P2A-VP1 VP0 VLPs, suggesting that the 2A peptide was degraded or cleaved by host factors on these VLPs but not those derived from VP3-P2A-VP1 VP0, indicating subtle differences in processing of these VLPs despite the similarity in production. This difference in particle composition may also account for the small differences seen in thermostability, as the VP3-P2A-VP0 VP1 VLPs appeared less thermally stable at 37°C, maintaining 62% D-antigenicity, whereas the VP3-P2A-VP1 VP0 VLPs maintained 77% D-antigenicity (Fig 5B.)

Importantly, the thermostabilities of the protease-independent VLPs were similar to those of the protease-dependent PV1 SC6b VLPs and maintained native antigenicity at physiologically relevant temperatures. These properties suggest that the protease-independent VLPs have the potential to induce protective immune responses against PV. However, the yields of protease-independent VLPs were lower than protease-dependent VLPs, likely due to inefficiency of the P2A translation re-initiation in *Pichia* as highlighted in Fig. 1B. This inefficiency of re-initiation may be addressable by manipulating the sequence surrounding the 2A peptide. In addition, the demonstration that native antigenic VLPs can be produced using protease-independent means may make these constructs more amenable from a genetic complexity perspective for adaptation to the new vaccine technologies, such as adenovirus-vectored vaccines and mRNA vaccines, that have emerged in response to the SARS-CoV-2 pandemic. They would also circumvent potential problems of cytotoxicity associated with 3CD and facilitate the production of immunogenic enterovirus VLPs *in vivo* (41–43).

In conclusion, we have shown protease-independent production of VLPs using a heterologous expression system, which maintain the antigenic, morphological and thermostability characteristics known to be important drivers of protective immunity against PV. Additionally, our data corroborate the results observed in other expression systems using this thermostable mutant, whilst building on this work to highlight that VP3 is the only structural protein able to tolerate the addition of a C-terminal P2A tag without negatively impacting assembly or antigenicity. Overall, the protease-independent VLP system we describe provides a framework for the production of VLPs using modern vaccine technologies, not only for PV but also as a model system for other members of the picornavirus family.

## Methods

### Vector construction

The P1 gene of PV1 SC6b Mahoney was amplified from a pT7RbzMahSC6bP1_deletion mutant plasmid sourced from NIBSC, UK and a 3CD gene was codon optimised for expression in *Pichia pastoris*. Both P1 genes and the 3CD were cloned separately into the pPink-HC expression vector multiple cloning site (MCS) using *EcoRI* and *FseI* (NEB). Subsequently, the dual promoter expression vector was constructed through PCR amplification from position 1 of the 3CD pPink-HC to position 1285 inserting a *SacII* restriction site at both the 5’ and 3’ end of the product. The P1 expression plasmids were linearised by *SacII* (NEB), followed by the insertion of the 3CD PCR product into *SacII*-linearized P1 plasmid. All PCR steps were carried out with Phusion polymerase (NEB) using the manufacturer’s guidelines. The PV1 SC6b 6xHIS P1 construct was subcloned by the addition of the 6xHIS tag through PCR amplification of the PV1 SC6b P1 from the PV1 SC6b P1 pPink-HC vector. The PV1 SC6b 6xHIS P1dual promoter expression vector was then constructed as described above. The PV1 SC6b P1 pPink-HC vector was used as the template to produce each porcine teschovirus 2A (P2A) containing construct. P2A was inserted into each construct through overlap PCR amplification and then dual promoter expression constructs for each P2A construct were obtained as described above for the 3CD containing plasmids.

### Yeast transformation and induction

Plasmids were linearized by *Afl*II digestion (NEB) and then transformed into PichiaPink™ Strain one (Invitrogen, USA) by electroporation as per the manufacturer’s guidelines. Transformed yeast cells were plated on *Pichia* Adenine Dropout (PAD) selection plates and incubated at 28°C until sufficient numbers of white colonies appeared (3-5 days). To screen for high-expression clones, 8 colonies were randomly selected for small-scale (5 mL) expression experiments. Briefly, colonies were cultured in YPD for 48 hours at 28°C and after shaking at 250 rpm, each culture was pelleted at 1500 × *g* and resuspended in YPM (1 mL & methanol 0.5% v/v) to induce protein expression and cultured for a further 48 hours. Cultures were fed methanol to 0.5% v/v 24 h post-induction. Expression levels of each clone were determined through VP1 expression analysed by immunoblotting as described below. For VLP production, a stab from a previously high-expressing glycerol stock was cultured for 48 hours in 5 mL YPD to high density. To increase biomass for the protease containing constructs, 4 mL of the starter culture was added to 200 mL YPD in a 2 L baffled flask and cultured at 28°C at 250 rpm for a further 24 h. Cells were pelleted at 1500 × *g* and resuspended in 200 mL YPM (methanol 0.5% v/v) and cultured for a further 48 h. Cultures were fed methanol to 0.5% v/v 24 h post-induction. For 2A containing constructs, the cultures were produced in the same way but in a total volume of 400 mL per construct. After 48 h cells were pelleted at 2000 × *g* and resuspended in breaking buffer (50 mM sodium phosphate, 5% glycerol, 1 mM EDTA, pH 7.4) and frozen prior to processing.

### Purification and concentration of PV and PV VLPs

*Pichia* cell suspensions were thawed and subjected to cell lysis using CF-1 cell disruptor at ~275 MPa chilled to 4°C following the addition of 0.1% Triton-X 100. The resulting lysate was then centrifuged at 5000 rpm to remove the larger cell debris, followed by a 10,000 × *g* spin to remove further insoluble material. The supernatant was then nuclease treated using 25 U/mL DENARASE® (c-LEcta) for 1.5 hours at RT with gentle agitation. The supernatant was then mixed with PEG 8000 (20% v/v) to a final concentration of 8% (v/v) and incubated at 4°C overnight. The precipitated protein was pelleted at 5,000 rpm and resuspended in PBS. The solution was then pelleted again at 5,000 rpm and the supernatant collected for a subsequent 10,000 × *g* spin to remove any insoluble material. The supernatant was collected and pelleted through a 30% (w/v) sucrose cushion at 151,000 × *g* (using a Beckman SW 32 Ti rotor) for 3.5 hours at 10°C. The resulting pellet was resuspended in PBS + NP-40 (1% v/v) + sodium deoxycholate (0.5% v/v) and clarified by centrifugation at 10,000 × *g*. Supernatant was purified through 15-45% (w/v) sucrose density gradient by ultracentrifugation at 151,000 × *g* (using a 17 mL Beckman SW32.1 Ti rotor) for 3 hours at 10°C (16). Gradients were collected in 1 mL fractions from top to bottom and analysed for the presence of VLPs through immunoblotting and ELISA. For electron microscopy and thermostability studies, peak fractions from primary gradients were diluted and purified through a second 15-45% (w/v) sucrose density gradient by ultracentrifugation at 151,000 xg for 3 hours at 10°C.

Peak gradient fractions as determined by immunoblotting and ELISA were then concentrated to ~100 uL in PBS + 20 mM EDTA using 0.5 mL 100 kDa centrifugal concentration columns (Amicon) as per the manufacturer’s instructions.

### Sample preparation and immunoblotting

Gradient fraction samples were mixed 5:1 with 5x Laemmli buffer and analysed by 12% SDS-PAGE (w/v) using standard protocols. Concentrated VLP samples were prepared at a 1:1 ratio using 2 x Laemmli buffer. Immunoblot analyses were performed using a monoclonal blend primary antibody against VP1 protein of each PV1, PV2, and PV3 (Millipore MAB8655) followed by detection with a goat anti-mouse secondary antibody conjugated to horseradish peroxidase, and developed using the chemiluminescent substrate (Promega). To detect VP0, a rabbit polyclonal antibody was used followed by detection with a goat anti-rabbit secondary antibody conjugated to horseradish peroxidase and developed using a chemiluminescent substrate (Promega). Immunoblot detection of 6xHIS and 2A peptide tags were determined through anti-Histidine tag (AD1.1.10, Biorad) and anti-2A peptide (3H4, Novus Biologicals), followed by detection with a goat anti-mouse secondary antibody conjugated to horseradish peroxidase, and developed using the chemiluminescent substrate (Promega) (44).

### Enzyme-linked immunosorbent assay (ELISA)

To determine the antigenic content of gradient fractions a non-competitive sandwich ELISA assay was used to measure PV1 D- and C-antigen content (45). Briefly, two-fold dilutions of antigen were captured using a PV1-specific polyclonal antibody, and then detected using anti-PV1 D-antigen (Mab 234) or C-antigen (Mab 1588) specific monoclonal antibodies (kindly provided by NIBSC), followed by anti-mouse peroxidase conjugate (46, 47). All ELISAs were then analysed through Biotek PowerWave XS2 plate reader.

### Thermostability Assay

The thermostability of VLPs was assessed using previously published protocols (32). Briefly, quantified PV VLPs were diluted in phosphate buffered saline (Corning 46-013-CM) to provide a uniform quantity of D antigen. Duplicate aliquots were incubated on ice (control) or in a thermocycler (BIO-RAD T100) at temperatures between 20 °C and 60 °C for 10 minutes.

Thermostability of the VLPs was assessed by measuring loss of D-antigenicity by ELISA, detection of D-antigenic particles was determined through PV-1 specific Mab 234.

### Electron microscopy

To prepare samples for negative stain transmission EM, carbon-coated 300-mesh copper grids were glow-discharged in air at 10 mA for 30 seconds. 3 μl aliquots of purified VLP stocks were applied to the grids for 30 seconds, then excess liquid was removed by blotting.

Grids were washed twice with 10 μl distilled H2O. Grids were stained with 10 μl 1% uranyl acetate solution, which was promptly removed by blotting before another application of 10 μl 1% uranyl acetate solution for 30 seconds. Grids were subsequently blotted to leave a thin film of stain, then air-dried. EM was performed using an FEI Tecnai G2-Spirit transmission electron microscope (operating at 120 kV with a field emission gun) with an Gatan Ultra Scan 4000 CCD camera (ABSL, University of Leeds).

### Image processing

Raw micrographs were visualised with ImageJ 1.51d (48, 49).

## Authors and Contributions

LS, DJR & NJS conceived and designed the experiments. LS, JJS, KG & HX conducted the experiments. LS, JJS, KG, DJR & NJS analysed the data. LS, DJR & NJS wrote the manuscript. KG, JJS, HX, MU & IJ reviewed and edited the manuscript. Funding was secured for this research by DJR and NJS.

## Acknowledgments

We thank other members of the Stonehouse/Rowlands group, at the University of Leeds, for their insightful contributions. This work was performed as part of a WHO-funded collaborative project involving the following institutions: University of Oxford, University of Reading, University of Florida, Harvard University, John Innes Centre, The Pirbright Institute, and the National Institute for Biological Standards and Control.

